# Coffee microsporogenesis and related small interfering RNAs biosynthesis have a unique pattern among eudicots suggesting a sensitivity to climate changes

**DOI:** 10.1101/2023.07.06.548025

**Authors:** Kellen Kauanne Pimenta de Oliveira, Raphael Ricon de Oliveira, Gabriel de Campos Rume, Thales Henrique Cherubino Ribeiro, Christiane Noronha Fernandes-Brum, Atul Kakrana, Sandra Mathioni, Blake C. Meyers, Matheus de Souza Gomes, Antonio Chalfun-Junior

## Abstract

Recently, the siRNAs pathways, and especially reproductive phasiRNAs, have attracted attention in eudicots since their biological roles are still unknown and their biogenesis took different evolutionary pathways compared to monocots. In this work, we used *Coffea arabica* L., a recently allotetraploid formed from the hybridization of *C. canephora* and *C. eugenioides* unreduced gametes, to explore microsporogenesis and small RNAs related pathways in a eudicot crop. First, we identified the microsporogenesis stages during anther development revealing that pre-meiosis occurs in anthers of 1.5 mm inside floral buds (FBs), whereas meiosis between 1.5 and 4.2 mm FBs, and post-meiosis in FBs larger than 4.2mm. These stages coincide with the Brazilian winter, a period of FBs reduced growth which suggests temperature sensitivity. Next, we identified and quantified the expression of reproductive 21- and 24-nt phasiRNAs during coffee anther development together with their canonical and novel miRNA triggers, and characterized the DCL and AGO families. Our results showed that the pattern of reproductive phasiRNA abundance in *C. arabica* is unique among described eudicots and the canonical trigger car-miR2275 is involved in the processing of both 21 and 24 nt phasiRNAs. Fourteen DCL genes were identified, but DCL5, related to phasiRNA biosynthesis in monocots, was not according to its specificity for monocots. Thus, our work explored the knowledge gap about microsporogenesis and related siRNAs pathways in coffee, contributing to the control of reproductive development and to the improvement of fertility in eudicots.

## 1 INTRODUCTION

Microsporogenesis is a fundamental process to produce pollen and guarantee fertilization of plants with sexual reproduction (Pradillo & Santos, 2018; Adhikari et al., 2020). This process is evolutionarily diverse in plants and largely governed by molecular pathways that include small interfering RNAs (siRNAs) and enzymes related to its biosynthesis (Ma et al., 2005; Albert et al., 2014; Åstrand et al., 2021). Recently, some studies have shed light on the function and biosynthesis of siRNA molecules, as phased secondary small interfering RNAs (phasiRNAs) and microRNAs (miRNAs) in monocot species (Nadot et al., 2008; Lee & Carroll, 2018; Dhaka et al., 2020). However, this knowledge is still poorly comprehended in eudicot species, especially crops with complex genomes and sexual reproduction cycles. Here, we explored this gap using *Coffea arabica* L. (Rubiaceae), a recent allotetraploid formed from the hybridization of *C. canephora* and *C. eugenioides* unreduced gametes (Lashermes et al., 1999; Scalabrin et al., 2020).

The phasiRNAs are small RNAs found in plants, generated from a long RNA precursor at intervals of 21 to 26 nucleotides (nt), hypothesized to be a relevant group for reproductive regulation (Xia et al., 2019; Zhai et al., 2015), these phasiRNAs are produced from non-coding precursors (“PHAS” transcripts) and transcribed from non-repetitive loci (*PHAS* loci) (Axtell, 2013a; Komiya, 2017). For example, reproductive phasiRNAs have a spatiotemporal pattern in monocots in which hundreds of *PHAS* loci on all chromosomes produce 21-nt phasiRNAs abundant in pre-meiotic anthers and 24 nt-phasiRNAs enriched in meiotic stage anthers (Fan et al., 2016; Kakrana et al., 2018; Zhai et al., 2015). Disturbance of 21-*PHAS* loci underlies photoperiod-sensitive male sterility in rice, while mutants that do not produce 24-nt phasiRNA produce conditional male sterility in maize (Zhai et al., 2015). Moreover, reproductive phasiRNAs are also present in female reproductive organs and therefore may function in both male and female germinal development (Kakrana et al., 2018).

The phasiRNAs can be classified into two main groups, based on their origin: from *PHAS* loci located in noncoding or protein-coding regions (Fei et al., 2013; Liu et al., 2020). Systematic analysis of the main phylogenetic plant groups, encompassing algae, mosses, gymnosperms, basal angiosperms, monocots, and eudicots, identify abundant *PHAS* loci among protein-coding genes, suggesting that phasiRNAs predominantly regulate these genes from which they are originate, in contrast to the dominant trans-regulatory mode of microRNAs (miRNAs) (Liu et al., 2020).

In monocots, two pathways associated with germlines are well described and produce diverse and abundant 21-nt (premeiotic) and 24-nt (meiotic) phased siRNAs (Fei et al., 2016; Zhai et al., 2015). These phasiRNAs are produced from capped and polyadenylated non-coding precursors (“PHAS” transcripts) and transcribed by RNA polymerase II (Pol II) from non-repetitive loci. They are typically triggered by two 22-nt miRNAs, miR2118/482 for 21-nt phasiRNAs and miR2275 for 24-nt phasiRNAs. 3’ mRNA fragments are converted to double-stranded RNA by RNA-DEPENDENT RNA POLYMERASE 6 (RDR6) and processed by DICER-like 4 (DCL4) and DICER-like 5 (DCL5) to produce 21-nt and 24-nt phasiRNAs, respectively. Then, phasiRNAs are loaded into ARGONAUTE (AGO) proteins to induce target RNA silencing (Kakrana et al., 2018; Komiya, 2017; Xia et al., 2019).

Notably, DCL5 is a Dicer protein found specifically in monocots and, so far, has not been reported in eudicots, despite the existence of 24-nt phasiRNAs and the conservation of miR2275, both broadly present in angiosperms (Xia et al., 2019). Moreover, genomic analyses showed divergence related to the miR2275/24-nt phasiRNA pathway in plants, since it is absent in legumes, *Arabidopsis,* and in *Solanaceae* species that produce 24-nt phasiRNAs but that lack the miR2275 trigger (Xia et al., 2019).

Evolutionary studies with non-grass monocots (garden asparagus, daylily, and lily) have suggested that the DCL5-dependent meiotic 24-nt phasiRNA pathway may have originated more than 117 MYA (KAKRANA et al., 2018). The DCL5 subfamily seems to have evolved from an ancient ‘eudicot-type’ DCL3 ancestor, able to produce 24-nt phasiRNAs, and DCL5 accumulated mutations specifying it for distinct substrates and ensuring the production of functional siRNAs (Chen et al., 2022). Thus, plants have diversified and optimized RNA silencing mechanisms to coordinate reproductive development during evolution (Chen et al., 2022). How functional differentiation of miRNA/DCL triggers and reproductive phasiRNAs was achieved, and their biological relevance, remains elusive, especially during the microsporogenesis of eudicot plants.

## 2 MATERIALS AND METHODS

### Plant material

Floral buds of *C. arabica* cv. Red Catuaí at different development stages (separated by length in millimeters) were collected in the experimental field of UFLA (−21.20549, −44.98021) to identify cells at pre-meiosis, meiosis, and post-meiosis. Tissues used for microscopy were fixed in Carnoy 3:1 and stored in the refrigerator at 4°C, whereas those used for RNA extractions were collected in liquid nitrogen and stored in an ultrafreezer at −80°C, until used.

### Optical and Scanning Electron Microscopy (SEM)

For optical microscopy, slides were prepared according to (Gardner; Rattenbury, 1974) protocol and pictures taken with a 400x magnifying glass with more than 4X digital zoom (Fig. 1A to C). For Scanning Electron Microscopy (SEM), floral buds were removed from the primary fixative and transferred to microtubes containing 0.05 M cacodylate buffer for ten minutes. The buffer was changed three times. Samples were immersed in 1% osmium tetroxide solution in a cacodylate buffer for two hours at room temperature. Finally, they were washed with distilled water three times and dehydrated in series with acetone (25%, 50%, 75%, 90% - 10 minutes each, and 100% three times for 10 minutes). After this step, samples were completely dried utilizing a sputtering apparatus (Balzers CPD 030). After drying, samples were mounted on stubs and submitted to gold metallization in the Balzers SCD 050 vaporizer and observed in SEM (LEO EVO 40 XVP (Carl Zeiss) with Bruker X-ray microanalysis system (Quantax EDS) and cryosystem (Gatan).

**Fig 1.**
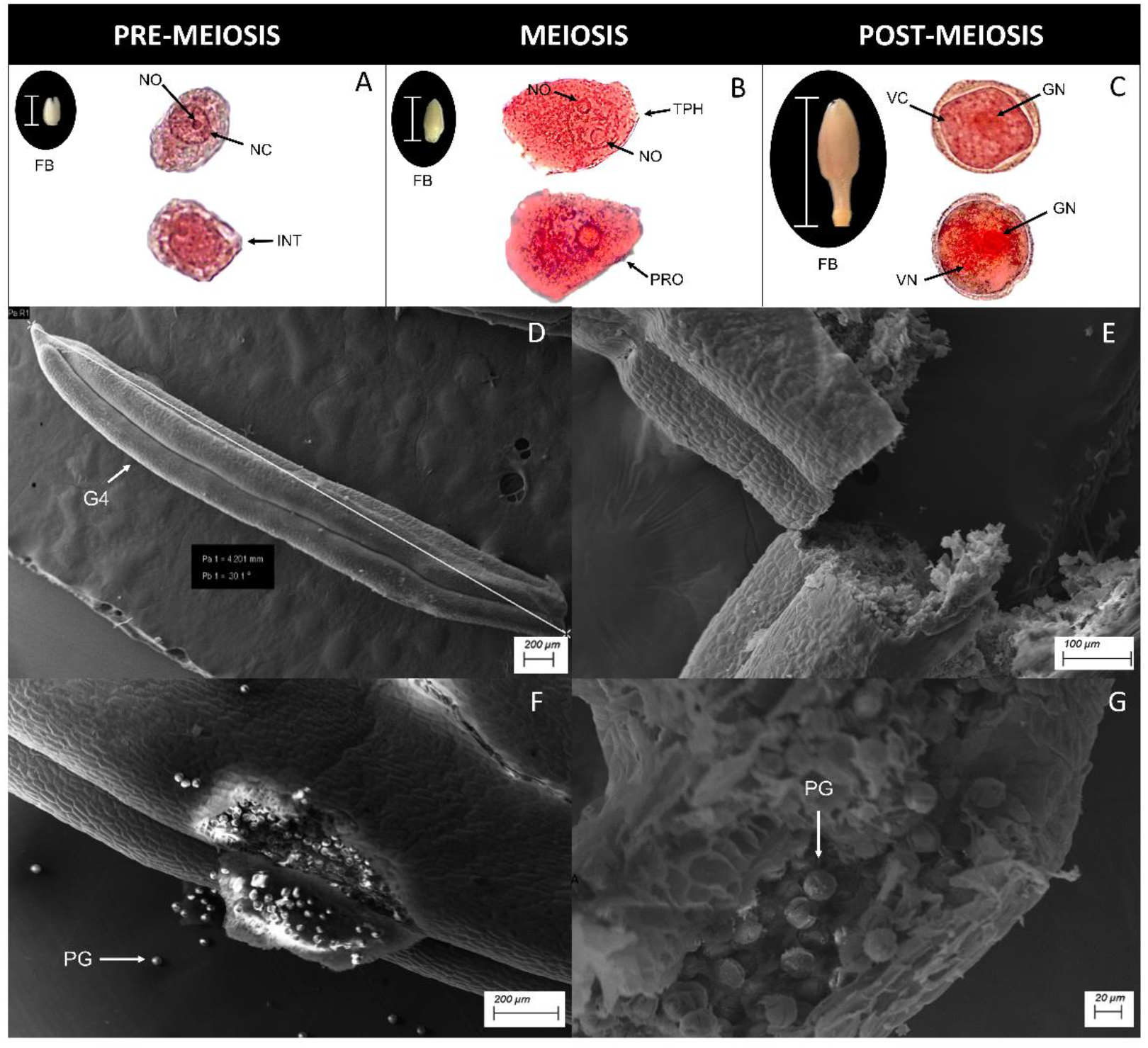
The microsporogenesis in *Coffea* sp.: cytogenetic analysis by optical and scanning electron microscopy (SEM) of coffee anthers at different developmental stages. Figure A to C show the pre-meiosis, meiosis and post-meiosis stages of microsporogenesis identified in coffee floral buds (FBs) with 1.5 mm, 4.0 mm and 15 mm, respectively. Figure D to G are pictures of SEM showing details of coffee anther and the released mature pollen. A) FB with 1.5 mm (bar in the figure) together with two one-cell microsporocytes at the pre-meiosis stage retrieved from the anthers. The microsporocytes are in the interphase (INT) with a well-defined nucleus (NC) and nucleolus (NO). B) FB with 4.0 mm (bar in the figure) together with two one-cell microsporocytes at the meiosis, one in prophase (PRO) I that marks the beginning of meiosis and other in telophase (TPH), with two visible nucleoli. C) FB with 15 mm (bar in the figure) together with two one-cell microsporocytes at the post-meiosis stage, one young vacuolated (VC) and mononucleated microspore, and one binucleated microspore with a generative nucleus (GN) and a vegetative nucleus (VN). D to G Pictures of anthers by SEM with mature pollen grains (PG) and size bars indicated in the figures.

### Small RNA libraries construction (Illumina sequencing)

Anthers from floral buds at different developmental stages were extracted and macerated in liquid nitrogen. Total RNA was extracted from 100 mg of tissue using the PureLink® RNA Mini Kit (Invitrogen) according to the manufacturer’s protocol. RNA samples were treated with DNase I using the RNase-free DNase Set (Qiagen) to eliminate residual DNA contamination. RNA integrity was visually analyzed in 1% agarose gel, and RNA content, as well as quality, were obtained by spectrophotometry (OD260/280 and OD260/230 > 1.8) (NanoVue GE Healthcare, Munich, Germany). RNA integrity number (RIN) was obtained with the Agilent 2100 Bioanalyzer system (Agilent Technologies) according to the manufacturer’s recommendations for plant RNA. Only samples with RIN greater than 7 were used in the following steps. Size selection for 20-30 nucleotide (nt) small RNAs was performed in denaturing Urea-PAGE gels, and libraries were constructed using the TruSeq Small RNA Library Preparation Kit (Illumina, cat # RS-200-0024) (Mathioni et al., 2017). All libraries were single-end sequenced with 51 cycles on an Illumina HiSeq 2500 instrument.

A total of 104.303.112 small-RNA-seq single-end reads were inspected for adapters with the Minion tool (Davis et al., 2013). The identified adapter sequences were removed with Trimmomatic v. 0.39 (Bolger et al., 2014). Only reads with more than 17 nucleotides were selected for further analysis after quality control. After sequencing and data cleanup steps, the small RNA libraries were aligned against *C. arabica* and *C. canephora* genomes databases, in which approximately 95% of the reads aligned in the first and 80% in the second. Details of the small RNA libraries of *C. arabica* anthers in pre-meiosis, meiosis and post-meiosis, including the numbers of Sequence Read Archive database (SRAs), are available in Table S1.

### Computational *de novo* identification of phased, siRNA-generating loci and their miRNA triggers

A total of 102.985.416 reads from pre-meiotic, meiotic, and post-meiotic small RNA libraries were submitted to the PHASIS suite (Kakrana et al., 2017). The first step was the de novo prediction of *PHAS* loci with the phase detect tool. Subsequently, phase merge was applied with parameter “-mode” set to “merge” to create a unified list of the PHASIS predicted in samples from the different stages. Finally, “phastrigs’’ were applied to identify miRNAs triggers for the previously identified *PHAS* loci; to do so we supplied to “phastrigs” a fasta file containing all the mature miRNAs identified in the “Conserved and novel miRNA prediction from *Coffea* sp” section. These steps were run twice to identify both 21 or 24-PHAS and the minimum significance p-value was set to 1e^-6^. Custom R-scripts were developed to plot heatmaps of the inferred abundance of each predicted PHAS.

### *In silico* and phylogenetic analysis of coffee DCL- and AGO-like proteins

To identify AGO and DCL homologs present in the coffee genome (available at National Center for Biotechnology Information’s (NCBI) webpage: https://www.ncbi.nlm.nih.gov/assembly/GCF_003713225.1/#/def) and perform phylogenetic analysis, we used the methods described in Rume et al., 2023. Briefly, sequences of AGO and DCL proteins of different species (see Tables S1) were downloaded from Genbank (Benson et al., 2009), UniProt (The Uniprot Consortium, 2021), and the MaizeGDB (Woodhouse et al., 2021). These sequences were utilized in separate alignments against the proteome of *C. arabica* using BLASTp v2.10.1 (Altschul et al., 1990) to retrieve similar sequences with an e-value of 10^-3^. To verify the presence of characteristic conserved domains (Fig. S2) from each protein family, candidate sequences were then analyzed using Conserved domains (https://www.ncbi.nlm.nih.gov/Structure/cdd/wrpsb.cgi), and filtered so that only one protein per gene locus remained. The phylogenetic analyses were made from the alignments of these identified coffee sequences together with those described in other species (Table S1).

### Transcriptional analysis of *DCLs* and *AGOs* in RNAseq libraries of different coffee tissues

To analyze the transcriptional abundance of the DCL and AGO genes identified, we used published and novel RNA-seq libraries from *C. arabica* (details in Table S1). Libraries were subjected to quality analyses using the program FastQC v.0.11.9 (Robinson et al., 2010) and trimmed as necessary with Trimmomatic v0.39 (Bolger et al., 2014). STAR v.2.7.8a (Dobin et al., 2013) with standard parameters was used to carry out fragment alignments against the genome of *C. arabica*. The resulting libraries were sorted, read duplicates were removed with the Picard toolkit (http://broadinstitute.github.io/picard), and mapped fragments were counted with the htseq-count algorithm (Anders et al., 2015). Expression values were calculated in log (RPKM + 1) using the edgeR package (Robinson et al., 2010), from the Bioconductor R v3.13 project (Huber et al., 2015).

### Conserved and novel miRNA prediction in *Coffea sp*

To search for putative conserved miRNAs and their precursors, we applied an adapted algorithm previously described by (De Souza Gomes et al., 2011) to the genome and transcriptome databases of *C. canephora* (Denoeud et al., 2014). The second prediction of conserved and novel miRNAs was made from RNA-seq libraries of *C. arabica* (de Oliveira et al., 2020; Cardon et al., 2022), in which mapped sRNA reads were used as input to two different computational pipelines for the discovery of miRNAs – a stringent pipeline for *de novo* identification and a relaxed pipeline for identification of conserved ‘known’ miRNAs (Jeong et al., 2013). Steps in both pipelines involved processing using Perl scripts as described earlier (Jeong et al., 2011), with a modified version of miREAP (https://sourceforge.net/projects/mireap/) and CentroidFold (Sato et al., 2009). In the ‘stringent’ criteria pipeline, sRNAs of length between 20 and 24 nt, with abundance ≥ 50 TP30M in at least one library, and total genome hits ≤ 20 were assessed for the potential pairing of miRNA and miRNA* using modified miREAP optimized for plant miRNA discovery with parameters –d 400 –f 25. Strand bias for precursors was computed as the ratio of all reads mapped to the sense strand against total reads mapped to both strands. In addition to strand bias, abundance bias was computed as the ratio of the two most abundant reads against all the reads mapped to the same precursor. Candidate precursors with strand bias ≥ 0.9 and abundance bias ≥ 0.7 were selected, and the foldback structure for the precursor was predicted using CentroidFold. Each precursor was manually inspected to match the criteria as described earlier (Jeong et al., 2013). All the miRNAs identified through this stringent pipeline were then annotated by matching mature sequences to miRBASE (version - 21), and those that did not match any known miRNA were considered lineage or species-specific. In the ‘relaxed’ criteria pipeline, which is implemented to maximize identification of ‘known’ miRNAs; relaxed filters were applied – sRNA between 20 and 24nt, with hits ≤ 20 and abundance ≥ 15 TP30M; and precursors with strand bias ≥ 0.7 and abundance bias ≥ 0.4. The stem-loop structure of candidate precursors was visually inspected, the same as the ‘stringent’ pipeline. Mature sequences of identified miRNAs were further matched with miRBASE entries (version-21), and those with total ‘variance’ (mismatches and overhangs) ≤ 4 were considered conserved miRNAs. Finally, sequences of the precursors of the two predictions were aligned and all parameters were compared. Redundant sequences were eliminated.

### Expression evaluation of conserved and novel miRNAs identified in the

### Coffea sp. genomes

A total of 102,985,416 reads from pre-meiotic, meiotic, and post-meiotic small-RNAs libraries were aligned against the conserved and putative novel miRNAs identified in the *Coffea sp*. The alignment was performed with ShortStack v. 3.8.5 (Axtell, 2013b) with the parameters “--mmap u --nohp -- locifile”. Custom R-scripts were developed to plot heatmaps of the inferred abundance of each putative novel miRNAs.

## 3 RESULTS

### Cytogenetic analysis identified the stages of microsporogenesis in *Coffea arabica*

Coffee anthers in different stages of development were harvested from floral buds (FBs) ranging from 1 to 15 millimeters and analyzed using optical microscopy and SEM. Cytogenetic analysis showed that microsporocytes in pre-meiosis I, with cells in interphase (INT), appear in anthers of floral buds up to 1.5 mm (Fig.1 A). In this stage, the nucleus (NC) and nucleolus (NO) are visible, reflecting the chromosome organization to duplicate and form chromatids (Conagin, 1961; Ma, 2005). The cells in meiosis were observed in anthers from floral buds of 1.5 to 4.2 mm (Fig. 1B). In this stage, it was possible to observe cells in two stages of meiosis, prophase (PRO) and telophase (TPH). In prophase, the chromosomes have begun to condense and the nucleoli and nuclear envelope are no longer visible. In the cells in telophase, the last phase of meiosis, the chromosomes unwound and it is possible to see two nucleoli and the nuclear envelope reappears (Conagin, 1961; Ma, 2005).

The post-meiosis stage was identified in anthers from floral buds larger than 4.2 mm, which presented mononucleated (VN) and vacuolated (VC) young microspores, with very thin cell walls, uniform, and clear cytoplasm, and a large, centralized nucleus (Fig. 1C). Binucleated microspores are also observed in post-meiosis, with a generative nucleus (GN) and a vegetative nucleus (VN). Soon after the post-meiotic mitosis, the generative nucleus can be seen attached to the microspore cell membrane; the nucleus then migrates to the center of the cell, already encased in its membrane system (Park & Twell, 2001). The external appearance of anthers in meiosis is shown in figure 1D, anthers with 4.2 mm or the G4 stage following Morais et al., (2008), in which no pollen grains are yet formed (Fig. 1E). The mature pollen occurs at the post-meiosis stage when anthers are larger than 4.2 mm and the tissue ruptures begin to appear (Fig. 1F and 1G).

### Identification and quantification of reproductive phasiRNAs and miRNAs in sRNA-seq libraries

In our analyses, we found seven 21-nt phasiRNAs and ten 24-nt phasiRNAs differently expressed in *C. arabica* sRNA-seq libraries corresponding to the microsporogenesis stages (Fig. 2 and Table S2). The 21-nt phasiRNAs were found to be more enriched in the post-meiotic stage while the 24nt-phasiRNAs are more abundant in the pre-meiotic stage (Fig. 5 = 2 and Table S3 => S2), in contrast to previous reports in monocots (Fan et al., 2016; Zhai et al., 2015). However, a recent study reported that 24-nt phasiRNAs representing more than 50% of all 24-PHAS annotated loci were enriched in pre-meiotic wheat anthers (Bélanger et al., 2020), which suggests that the pattern of accumulation of reproductive phasiRNAs varies across species. The meiotic stage has a lower expression of phasiRNAs in both monocots and eudicots.

**Fig 2.**
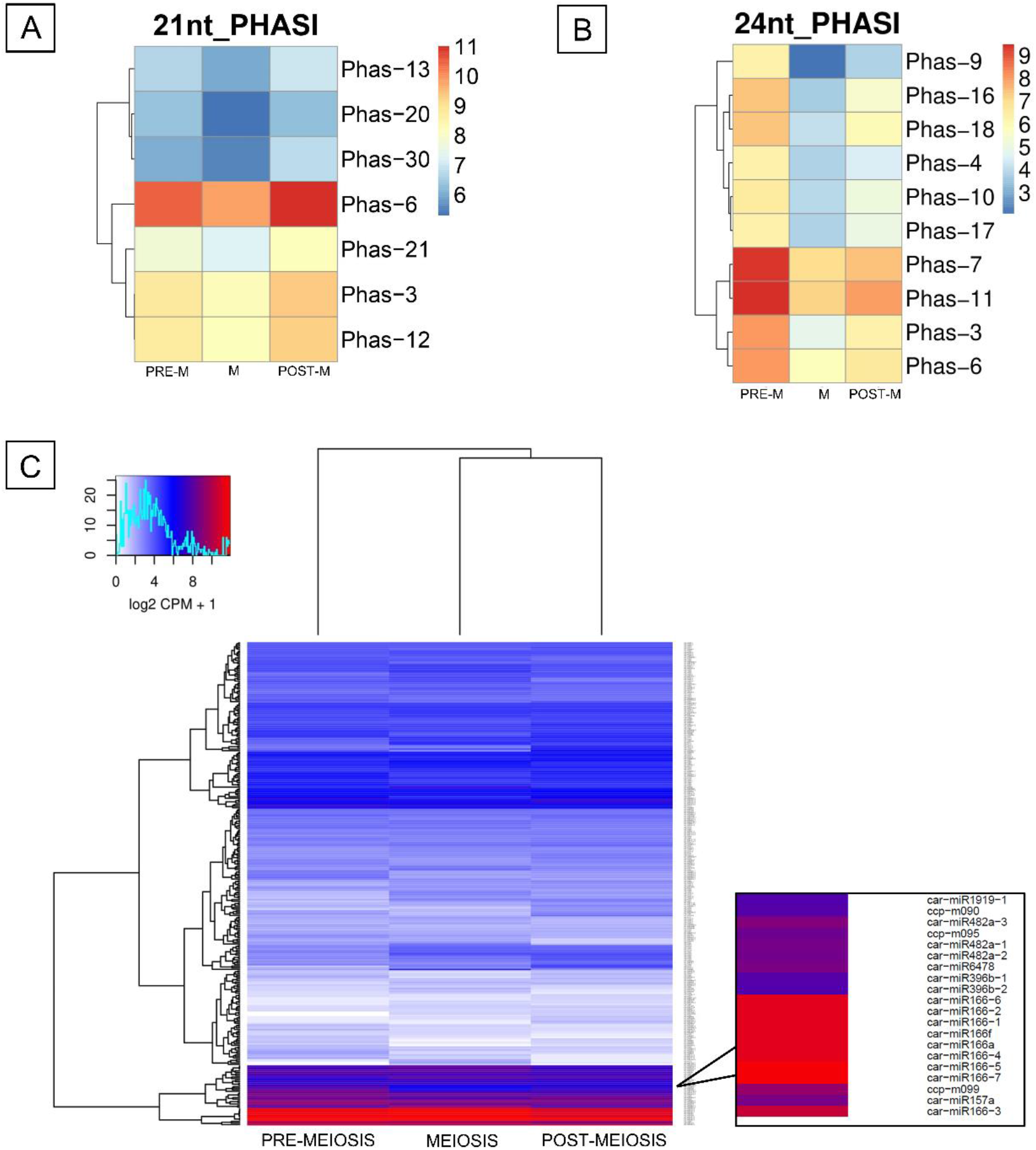
Expression profile of phasi and miRNAs identified in *Coffea arabica* L. A: Heatmap of 21-nt phasiRNAs (lines) identified in libraries (columns) of anthers in pre-meiosis (PRE-M), meiosis (M), and post-meiosis (POST-M). B: Heatmap of 24-nt phasiRNAs (lines) identified in libraries (columns) of anthers in pre-meiosis (PRE-M), meiosis (M), and post (POST-M). C: Heatmap of conserved and novel miRNAs (lines) identified in libraries (columns) of anthers in pre-meiosis, meiosis, and post-meiosis. The dendrogram on top of the heatmaps reflects the relationship of different conditions while the dendrogram on the left reflects the relationship of different sRNAs across conditions. In figures A and B, the colors are coded as a function of logarithmic base 10 of expression values normalized in FPKM +1 (Fragments per million mapped reads), whereas in figure C colors are coded as a function of logarithmic base 2 of expression values normalized in CPM+1 (Counts per million mapped reads).

miR2118/482 is described as responsible for processing 21-nt phasiRNAs while miR2275 processes 24-nt phasiRNAs (Kakrana et al., 2018; Komiya, 2017; Xia et al., 2019). Thus, to evaluate the presence of these miRNA triggers and others probably related to microsporogenesis development, we identified the miRNA precursors in the *Coffea canephora* genome (Table S3). From this, 138 miRNA precursors and 149 gender-specific mature miRNAs, 80 of which were miRNA-5p and 69 miRNA-3p, were identified in the *Coffea canephora* genome (Table S3). We also identified 259 conserved miRNA precursors that gave rise to 349 mature miRNAs, including 98 miRNA-5p and 251 miRNA-3p sequences (Table S3).

The main miRNA families were found together with conserved and lineage-specific miRNAs, and were quantified in the sRNA libraries; for example, miR157, miR396, and miR482 families were some of the most abundant in all the stages of microsporogenesis in *C. arabica* (Fig. 2C). The miR166 was also found to be notably abundant (Fig. 2C), this abundance of miR166 in Illumina is a common phenomenon, as noted in Baldrich, et al. (2015). In our results, car-miR482 is one of the main triggers for 21-nt phasiRNAs, whereas car-miR2275 appears as a trigger for car-miR2275 both 21-nt and 24-nt phasiRNAs (Table S4). We have not identified miR2118 in our data, so in addition to car-miR2275, results suggest that other miRNAs can cleave 24-nt phasiRNAs.

### Characterization of DCL and AGO proteins in the *Coffea arabica* genome

DICER-like (DCL) and ARGONAUTE proteins (AGO) are two families of proteins related to the production of siRNAs and phasiRNAs, which are generated by specific and divergent members in plants (Baumberger & Baulcombe, 2005; Liu et al., 2009). Thus, to identify DCLs and AGOs and the possibility to generate phasiRNAs during microsporogenesis, we characterized the two families in the *C. arabica* genome. As a result, we found fourteen DCL and twenty AGO coffee proteins that we further classified through phylogenetic analyses (Figs. 3 and 4), comparing them with homologous sequences of other monocot and eudicot plants (Table S1). We found three copies of CaDCL1, in which CaDCL1.1 and CaDCL1.2 were highly conserved and considered the same, but derived from a different parental subgenome, while CaDCL1.3 was truncated (Table S1 and Fig. S1); The DCL2 clade was the group with the highest number of CaDCLs homologs, eight, but only CaDCL2.1 presented the complete main domains (Table S1 and Fig. S1); two DCL3 copies were found to be similar in sequence but from different parental subgenomes; and finally, only one CaDCL4 copy was found, from the *C. eugenioides*’ subgenome (Table S1). Compared to other works, here we found five more DCLs than previously described by Mosharaf et al. (2019) and eight more than those published by Noronha Fernandes-Brum et al. (2017). Details of all the sequences are available in Table S1.

**Fig 3.**
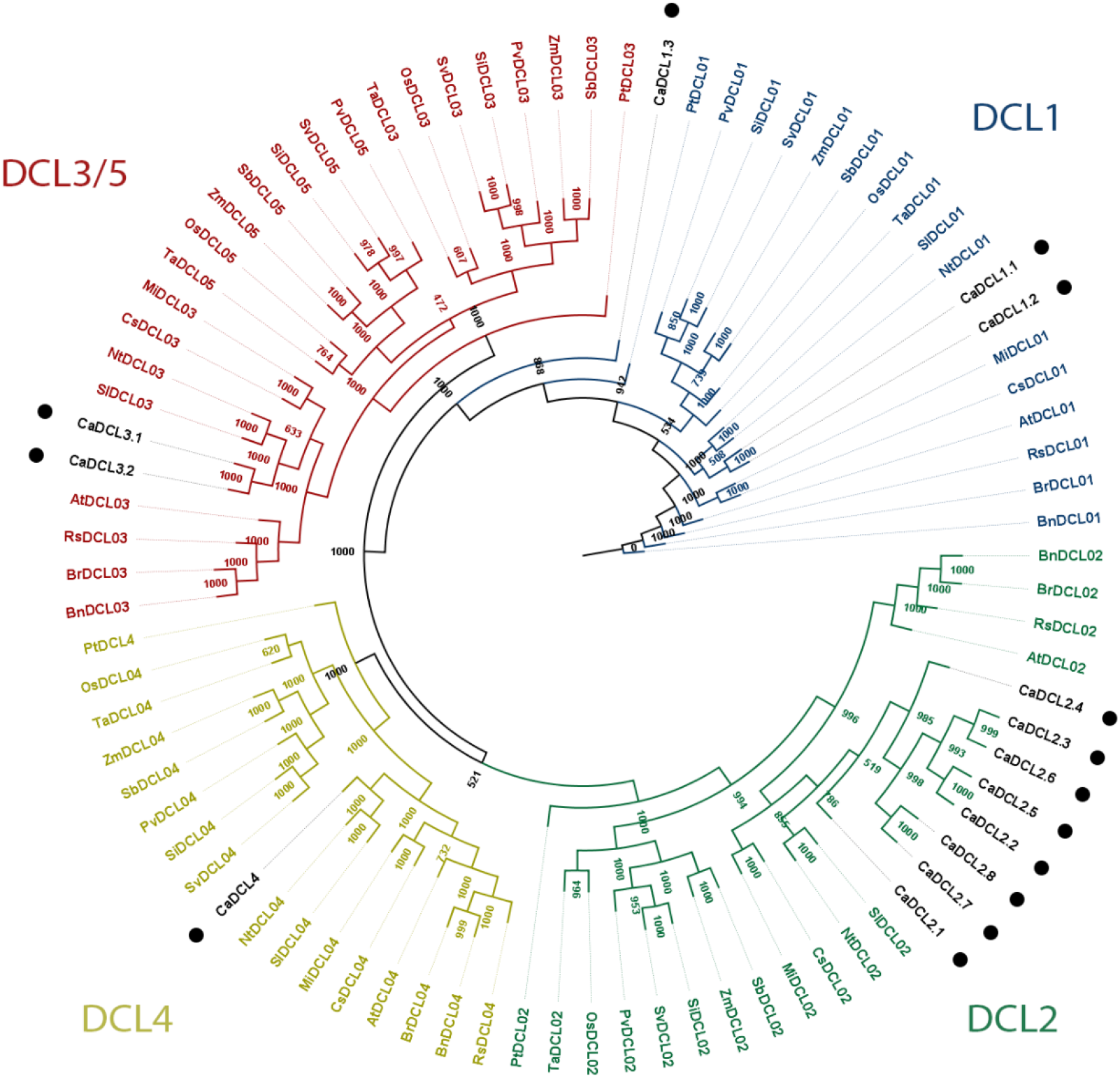
Phylogenetic analysis of coffee Dicer-like (DCL) proteins. The tree was generated through the neighbor-joining method containing DCL protein sequences identified in *C. arabica* (black dots) together with sequences of monocots, eudicots and gymnosperms. Details of all the sequences used are described in Table S1. The four major clades of DCLs were retrieved from sequences and indicated around the tree, being the group DCL3/DCL5 joined with DCL5, found only in monocots, due its evolutionary relationship (CHEN et al., 2022). A total of fourteen *C. arabica* DCL sequences were identified, three DCL1 homologs, eight DCL2, two DCL3 and one DCL4 (six complete sequences and eight truncated: CaDCL1.3, CaDCL2.2, CaDCL2.3, CaDCL2.4, CaDCL2.5, CaDCL2.6, CaDCL2.7, CaDCL2.8). Legends: *Arabidopsis thaliana* (At), Mangifera indica (Mi), Brassica napus (Bn), Raphanus sativus (Rs), Citrus sinensis (Cs), Brassica rapa (Br), Nicotiana tabacum (Nt), Solanum lycopersicum (Sl), Vitis vinifera (Vv), Fragaria vesca (Fv), Zea mays (Zm), Sorghum bicolor (Sb), Setaria viridis (Sv), Setaria italica (Si), Panicum virgatum (Pv), Oryza sativa (Os), Triticum aestivum (Ta), Pinus tabuliformis (Pt).

**Fig 4.**
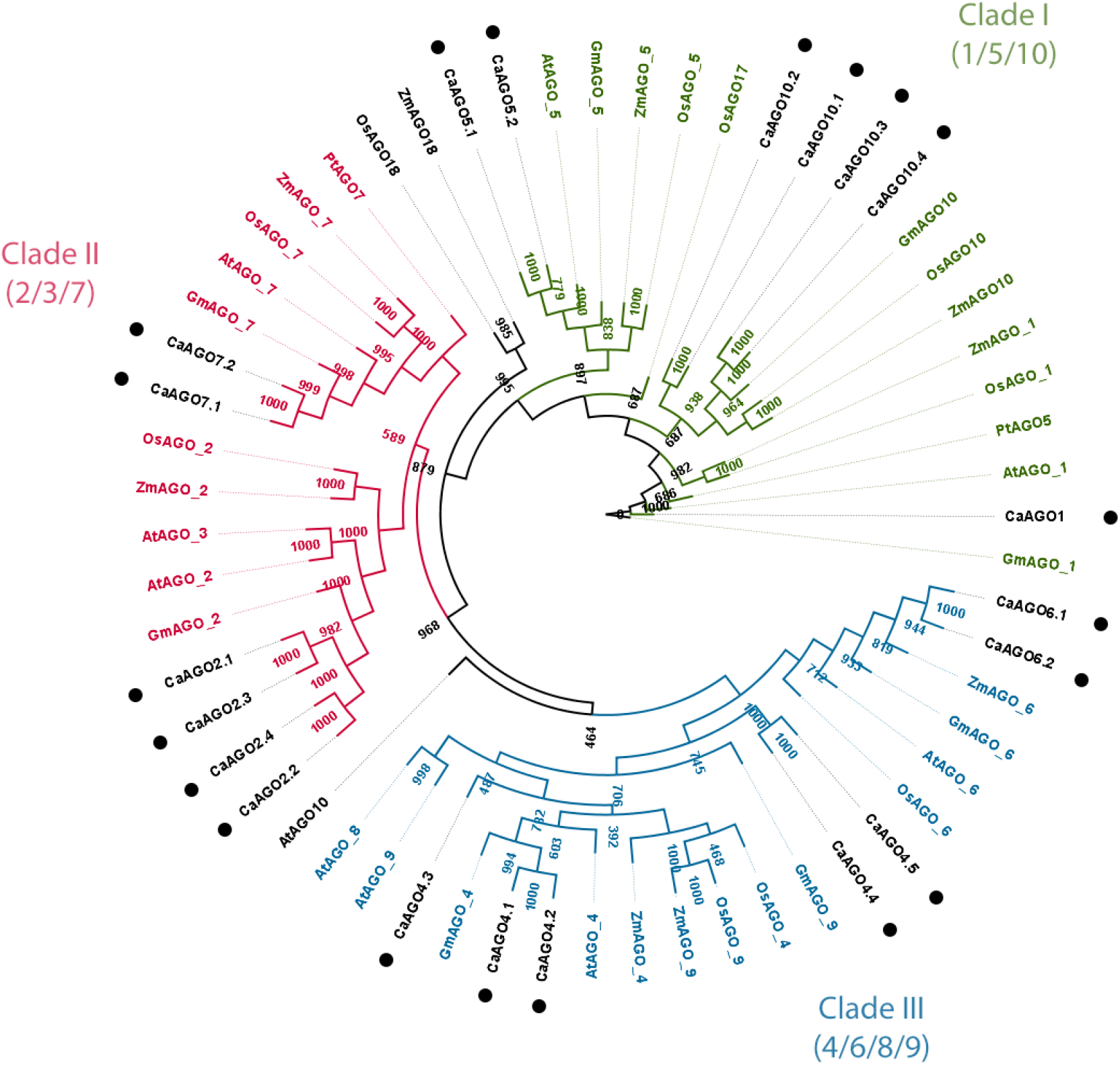
Phylogenetic analysis of coffee ARGONAUTA (AGO) proteins. The tree was generated through the neighbor-joining method containing AGO protein sequences identified in *C. arabica* (black dots) together with sequences of monocots, eudicots and gymnosperms. Details of all the sequences used are described in Table S1. The three major clades of AGOs were retrieved from sequences and indicated around the tree, clade I (AGO1/5/10), clade II (AGO2/3/7) and clade III (AGO4/6/8/9). A total of twenty *C. arabica* AGO sequences were identified, one AGO1 homolog, four AGO2, five AGO5, two AGO6, two AGO7 and ten AGO10. Legends: *Arabidopsis thaliana* (At), Glycine max (Gm), Oryza sativa (Os), Zea mays (Zm), Pinus tabuliformis (Pt).

Dicer and DCL proteins are a unique class of ribonucleases that generally contain six types of domains, including the DEAD-box, helicase-C, DUF283, PAZ, RNase III, and dsRBD domains. However, for some species, one or more of these domains may be absent (Liu et al., 2009; Margis et al., 2006). All identified CaDCLs presented Ribonuclease III domains, which have been considered one of the key domains for DCL proteins (Nicholson, 2014), except for CaDCL1.3 which only presented two (Fig. S1). In addition, CaDCL1.1, CaDCL1.2, CaDCL2.1, CaDCL3.1, CaDCL3.2, and CaDCL4 had the highest number of domains in common, Ribonuclease III, DEAD/RES III, Helicase_C, Dicer_dimer, PAZ, and DSRM (Fig. S1). Last instead being a divergent double-stranded RNA-binding domain coinciding with the DUF283 of Dicer (Dlakić, 2006).

Flowering plants have ten or more AGO proteins (Zhai et al., 2015). In total, 20 AGO-like proteins were found in the genome of *C. arabica* and distributed in three distinct clades of the phylogenetic tree (Fig 4), one CaAGO1, four CaAGO2, five CaAGO4, two CaAGO5, two CaAGO6, two CaAGO7, and four CaAGO10 were identified (Table S1). CaAGO1, CaAGO5, and CaAGO10 proteins were clustered into clade 1 which comprises the AGO1/5/10 proteins (Fig 4). Proteins CaAGO2 and CaAGO7 were grouped into clade II, which comprises AGO2/3/7 proteins. CaAGO4 and CaAGO 6 proteins were clustered into clade III, which comprises the AGO4/6/8/9 proteins. Among the AGO family members identified, all had copies in both subgenomes of *C. arabica*, except for CaAGO1, for which we identified only one copy on chromosome 4 from *C. canephora* (Table S1). In addition to the AGO sequences previously identified in the genome of *C. canephora* (Noronha Fernandes-Brum et al., 2017), we identified two more copies of AGO2, one copy of AGO5, and three AGO4 from *C. eugenioides*’ chromosomes. We also identified one new AGO6 and two new copies of AGO10, one from each of the two subgenomes.

AGO proteins are characterized by ArgoN (amino-terminal N), PAZ (PIWI– ARGONAUTE–ZWILLE), MID (middle), and PIWI domains (Meister, 2013). The conserved domains of the CaAGO proteins were analyzed, and all of them presented ArgoN, PAZ, PIWI, and ArgoL1 domains (Fig. S2). CaAGO1, CaAGO4.1, CaAGO4.2, CaAGO10.1, CaAGO10.2, CaAGO10.3, and CaAGO10.4 had all 6 domains ArgoN, PAZ, Piwi, ArgoL1, ArgoL2 and ArgoMID. The AGO1 proteins have an additional N-glycine-rich region (Gly-rich Ago1) domain that was present both in the CaAGO1 identified in this work and previously (in the CcAGO1; Fernandes-Brum et al. 2017). The MID domain is present in eight identified CaAGOs (CaAGO1, CaAGO 4.1, CaAGO 4.2, CaAGO 5.1, CaAGO 10.1, CaAGO 10.2, CaAGO 10.3, CaAGO 10.4). The MID domain anchors the 5′ ends of the small RNA by providing a binding pocket in which the 5′ terminal base engages in stacking interactions with a conserved tyrosine. In addition, several hydrogen bonds coordinate correct 5′ end binding (Meister, 2013).

### Expression analyses of DCL and AGO genes in different RNAseq libraries

To analyze the expression profile of CaDCLs and CaAGOs identified in this work in different tissues of the coffee tree, we aligned these sequences against RNA-seq libraries from *C. arabica* tissues under different conditions: leaves from plants under optimal and warm temperature conditions (cv. Acauã and Catuaí); FBs at G4 and G5 stages (cv. Catuaí and Siriema), embryos, stems, roots, meristems, floral buds, green drupes, yellow drupes, red drupes, fruits at multiple stages of development, fruits 30 days after flowering, and fruits 90 days after flowering (see details in Table S1).

All fourteen CaDCLs identified here were found to be expressed in at least one of these libraries (Fig. 5A). CaDCL2.1, CaDCL2.2, CaDCL3.1, CaDCL3.2 and CaDCL4 were the most highly expressed, with CaDCL3.2 more expressed in the reproductive libraries that correspond to FBs with 3.1 to 10 mm or the G4 and G5 stages described by Morais et al. (2008; G4_CA, G5_CA, G4_SE, and G5_SE in Fig. 5A). Of the twenty AGO proteins identified in this work, eighteen were expressed in the analyzed libraries (Fig. 5B). *CaAGO1* was the most expressed and appeared in all libraries, whereas CaAGO4.4 and CaAGO4.5 were not found expressed in any of the libraries.

**Fig 5.**
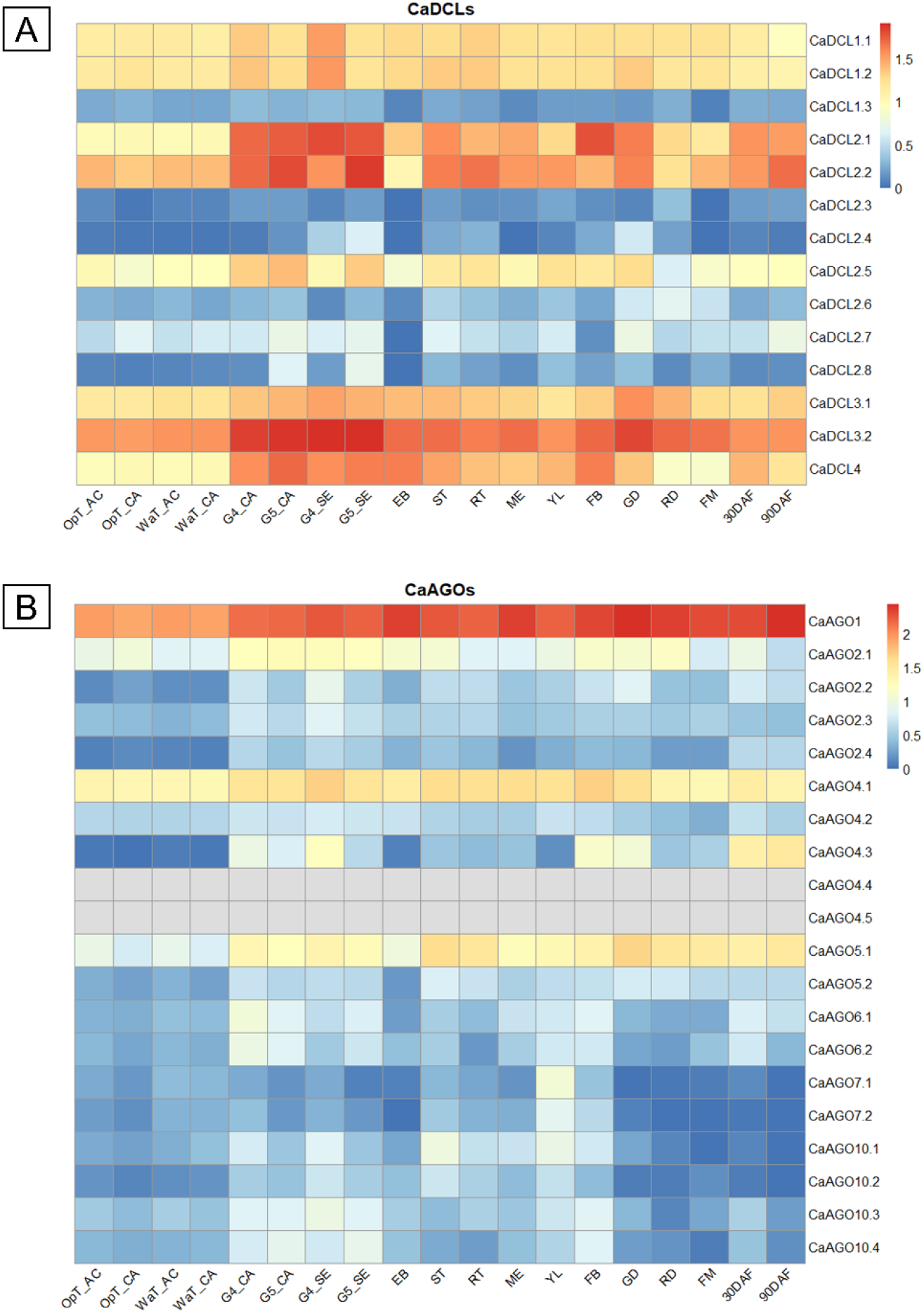
Expression profile of DCLs (A) and AGOs (B) genes identified in *Coffea arabica* L. The coffee genes are indicated in the right column of each figure and the RNA-Seq libraries relative to different coffee tissues, in the bottom line. Colors are coded as a function of logarithmic base 10 of expression values normalized in FPKM +1 (Fragments per million mapped reads). After validation of RNA-Seq quality parameters (details in Table S1 and Material and Methods), the expression profiles were determined in 19 different coffee tissues: OpT_AC and OpT_CA, *C. arabica* cv. Acauã (AC) and Catuaí (CA) leaves from plants in optimal temperature conditions; WaT_AC and WaT_CA, *C. arabica* cv. Acauã (AC) and Catuaí (CA) leaves from plants in warm temperature conditions; G4_CA and SE: floral buds at G4 stage from *C. arabica* cv. Catuaí (CA) and Siriema (SE); G5_CA and SE: floral buds at G4 stage from *C. arabica* cv. Catuaí (CA) and Siriema (SE); EB: embryos; ST: stems; RT: roots; ME: meristems; YL: yellow drupes; FB: floral buds; GD: green drupes; RD: red drupes; FM: fruits at multiple stages of development; 30DAF: fruits 30 days after flowering; 90DAF: fruits 90 days after flowering. The accession number of these RNAseq libraries and references are available at Table S1.

## 4 DISCUSSION

To comprehend the microsporogenesis and associated molecular pathways governing anther development in coffee, the cytological stages of meiosis were first determined. Here, we identified that the crucial steps of microsporogenesis, that is, pre-meiosis, meiosis, and post-meiosis, occur in FBs of length, respectively, less than 1.5 mm, 1.5 to 4.2 mm, and greater than 4.2 mm (Fig. 1A to C). Mature pollen grains are only observed after these stages (Fig. 1D to G). Compared to established classifications of coffee floral development, these meiosis stages are recognized in the G3 and G4 stages following Morais et al. (2008) and in Stage 4 following De Oliveira et al. (2014). Interestingly, coffee meiosis coincides with the Brazilian winter, marked by drought and low temperatures, a period of reduced growth of FBs, interpreted as dormant or latent (reviewed by López et al., 2021). A discussion about this terminology is provided by Considine & Considine (2016), but here we adopt the term “latent” because there is no evidence of the physiological basis of coffee dormancy (López et al. (2021)

Thus, here we show that the coffee meiosis occurs when FBs are apparently “dormant or latent” in the G3/G4 stage, since no external growth is visible but inside there is a high activity at the cellular level due to gametogenesis. This agrees with the reported post-meiosis observed in anthers of FBs with 6 to 8 mm, that contains microspores shortly after tetrad formation with the majority in mid-to late-uninucleate stage (Neuenschwander & Baumann, 1995; De Oliveira et al., 2014). Considering that microspore mitosis took place two to three days before anthesis in FBs of length 12-15 mm (Neuenschwander & Baumann, 1995), we now have the completed cycle of microsporogenesis and anthers development described.

Since the external growth at G3/G4 stages decreases, while an intense gametic development occurs within FBs, it is tempting to speculate that the plant’s metabolic energy is shifted to gametogenesis. Moreover, it seems plausible to associate this energetic control to temperature sensitivity during gametogenesis. Coffee is sensitive to warmth, changing its sugar content (De Oliveira et al., 2020), and some studies have reported that lower temperatures dramatically enhance pollen vigor while higher temperatures have the opposite effect (Wang et al., 2023). In accordance, pre-culturing FBs for two days at 4 °C derange the first division of the microspore nucleus (Neuenschwander & Baumann, 1995). Thus, while cold seems to have a beneficial role in coffee gametogenesis, the opposite seems to be true, with warming producing unreduced gametes, as observed in other species (Pécrix et al. 2011; Wang et al., 2017). Since a high temperature environment has the potential to increase gamete ploidy levels (Pécrix et al. 2011), this could explain the formation of diplogametes and the allotetraploid of *C. arabica* that originated in a period of global adverse conditions. Exploring these hypotheses and associated molecular mechanisms is necessary to make advances in the control of coffee fertility and ploidy for plant breeding programs.

Reproductive phasiRNAs are crucial for the microsporogenesis of many species, especially monocots (Liu & Wang, 2021; Liu et al., 2020). It was previously believed that only this group had 24-nt phasiRNAs, because their processing depends on a specialized DCL5 never reported in eudicots. However, recent studies have demonstrated the presence of these phasiRNAs in eudicots (Xia et al., 2019) and now we have also identified this class of molecules in *C. arabica*. Compared to monocots and other eudicots, we identified a smaller number of reproductive phasiRNAs, and a specific temporal pattern, with the 21 nt-phasiRNAs being more expressed in post-meiosis, and not in pre-meiosis, and the 24-nt phasiRNAs in pre-meiosis and not in meiosis (Kakrana et al., 2018; Pokhrel et al.,, 2021; Xia et al., 2019), consistent with the recent report in wheat by Bélanger, et al. (2020).

Another relevant element in the biogenesis of reproductive phasiRNAs is the miRNAs that act as their triggers. In several species, such as barley and wheat (Bélanger et al., 2020), and members of the family *Solanaceae* (Xia et al., 2019b), other triggers for 24-nt phasiRNAs have been described for in addition to the canonical miRNA miR2275. Interestingly, miR2118/482, known to trigger 21-nt phasiRNAs in eudicots and monocots, also has a role in generating 24-nt phasiRNAs in the eudicots columbine and wild strawberry. (Pokhrel et al., 2021). This suggests that miR2118/482 members present in gymnosperms may initiate 24-nt phasiRNA expression in angiosperms/basal eudicots, but miR2275 may have arisen- and therefore assumed this role-later. In *C. arabica*, miR2118 seems to have been lost, while the miR482 family acts as a trigger for 21-nt phasiRNAs, possibly together with the miR2275 family that is supposed to have assumed a role in the processing of both reproductive phasiRNAs. Further experiments are necessary to investigate this hypothesis.

The recent discovery of 24-nt phasiRNAas in eudicots drew attention to which DCL could be involved in their biosynthesis process since there is no DCL5 reported yet (Liu et al., 2020; Pokhrel et al., 2021; Xia et al., 2019). The emergence of DCL5 in monocots is believed to be explained by a subfunctionalization of the ancestral DCL3 protein, which is suggested to function in the production of heterochromatic siRNAs (hc-siRNAs) and phasiRNAs (Liu et al., 2020). In rice, OsDCL3 and OsDCL5 have completely different substrate specificities, while the eudicot AtDCL3 has an intermediate preference for dsRNAs with an overhanging 5’ triphosphate and 3’ structure (Chen et al., 2022). This suggests that monocot DCL5 and DCL3 proteins were optimized for cognate substrates after duplication of the progenitor ‘eudicot-like’ DCL3 protein and this functional specialization appears to have been achieved through the accumulation of mutations in the PAZ domain (Chen et al., 2022). Therefore, it is hypothesized that DCL3 has dual functions in some eudicots, processing distinct substrates (Pol II vs Pol IV RNA) for the biogenesis of 24-nt phasiRNAs or 24-nt hc-siRNAs (Xia et al., 2019). Our results support this (Fig. 3 and S1), and future work testing protein activity should explore this possibility.

Interestingly, of the three *CaDCL* genes expressed in the tissues analyzed in this study, *CaDCL3.2* was more abundant in the libraries of FBs in the G4 and G5 stages (Fig. 5A). Such stages correspond to the meiosis and post-meiosis stages identified here, supporting the hypothesis of a DCL3 role in the biogenesis of siRNAs during coffee microsporogenesis. DCL4, described as responsible for cleaving 21-nt phasiRNAs, identified in this work, was expressed in all analyzed tissues, mainly in G4, G5 and embryos, which reinforces the involvement of this molecule in the process (Fig. 5A) (Kakrana et al., 2018; Komiya, 2017; Xia et al., 2019). Each class of DCL has evolved to participate in a primary pathway, however, DCL2/DCL4 can also function as partial overlap, because defects in one class of DCL may be compensated by other classes in some cases (Fukudome & Fukuhara, 2017; Pokhrel et al., 2021).

In our study we identified twenty members of the CaAGO-like protein family (Fig. 4). The high number of AGO protein family members in flowering plants points to functional diversification, presumably reflecting regulatory pathways targeting the expanding sRNA class of molecules (Zhang et al., 2015). Members of the AGO4/6/8/9 clade have been described as effectors of 24 nt hc-siRNAs and are known to function in the regulation of transcription of genes (Zhang et al. 2015). In some eudicots, studies based on phylogenetic and expression data suggested that AGO5 and AGO1 could act as effector proteins for 21-nt phasiRNAs and AGO6 and AGO9 for 24-nt phasiRNAs (Pokhrel et al., 2021). In our data, at least one copy of each member of the *CaAGO*-like gene family was expressed in the libraries (Fig. 5B), with CaAGO4.2 and CaAGO4.3 being more abundant in the reproductive libraries, suggesting the role of the codified enzymes in the biogenesis of 24-nt phasiRNAs. *CaAGO6* was not expressed in the analyzed libraries and we did not identify an AGO9 homolog in our data.

## 5 CONCLUSIONS

In this work, we used *Coffea arabica* L., a recent allotetraploid formed from the hybridization of *C. canephora* and *C. eugenioides* unreduced gametes, to explore microsporogenesis and small RNA related pathways in a eudicot crop. We identified the pre-meiosis, meiosis and post-meiosis stages during anther development and, together with other work, provided a more complete overview of coffee microsporogenesis. Such stages coincide with the cold period in Brazilian conditions, which was associated with a benefit for pollen formation; on the contrary, warm conditions could be related to the formation of unreduced gametes and ploidy disturbance. Next, to explore siRNAs pathways, we identified and quantified the expression of reproductive 21-and 24-nt phasiRNAs, related miRNAs, and DLCs and AGOs. The presence of reproductive phasiRNAs in *C. arabica* presents a temporal pattern different from most of the eudicots already studied, with 21-phasi RNAs enriched in post-meiosis and 24-nt phasi-RNAs in pre-meiosis, which could reflect a diversification along the evolutionary history of woody eudicot species. The triggers identified here reinforce data that canonical microRNAs such as miR2275 may be involved in the processing of both 21 and 24-nt reproductive phasiRNAs. Despite fourteen DCLs being found here, DCL5 was not found in coffee; DCL5 is related to phasiRNA biosynthesis in monocots, and its absence here supports the hypothesis that this function could have evolved and been maintained by a closely related DCL3-like protein. Thus, our work contributes to highlighting the microsporogenesis and related reproductive siRNAs pathways in coffee and contributes towards the control of reproductive development and improvement of fertility in eudicots of economic interest.

## AUTHORS’ CONTRIBUTIONS

R.R.O., C.N.F.B., and A.C-J. conceptualized the project; K.K.P.O. conducted all data research and analyses; G.C.R., T.H.C.R., A.K., S.M, M.S.G. and B.C.M. supported for bioinformatic analyses; R.R.O. and A.C-J supervised experiments and analyses. K.K.P.O and R.R.O. wrote the manuscript; all co-authors corrected and contributed to writing of the final manuscript version.

## ACKNOWLEDGEMENTS

We thank the Federal University of Lavras (UFLA/Brazil) and members of the Laboratory of Plant Molecular Physiology (LFMP, UFLA/Brazil) for structural support of the experiments and analysis. We are grateful to Dr. Joanna Friesner for her helpful comments and editing of the manuscript. We also thank professor Vânia Helena Techio of the Biology Institute (UFLA/Brazil) for technical support in the cytogenetic analyses. Finally, we thank Instituto Brasileiro de Ciência e Tecnologia do Café (INCT/Café), Coordenação de Aperfeiçoamento de Pessoal de Nível Superior (CAPES), Conselho Nacional de Desenvolvimento Científico e Tecnológico (CNPq) and Fundação de Amparo à Pesquisa de Minas Gerais (FAPEMIG) for the financial support.

## DATA AVAILABILITY

Main data supporting the findings of this study are available within the paper and within its supplementary materials published online. The raw data used for analyses and figures are available from the corresponding author, Antonio Chalfun-Júnior, upon request. Raw data for the RNAseq runs can be obtained through BioProject ID PRJNA923233. Raw data for the smallRNAseq runs can be obtained through BioProject ID PRJNA925088. The respective SRA accessions are available in Supplemental Table S1.

