## Supplementary Figures for "Coffee microsporogenesis and related small interfering RNAs biosynthesis have a unique pattern among eudicots suggesting a sensitivity to climate changes"

**Supplementary Figures**
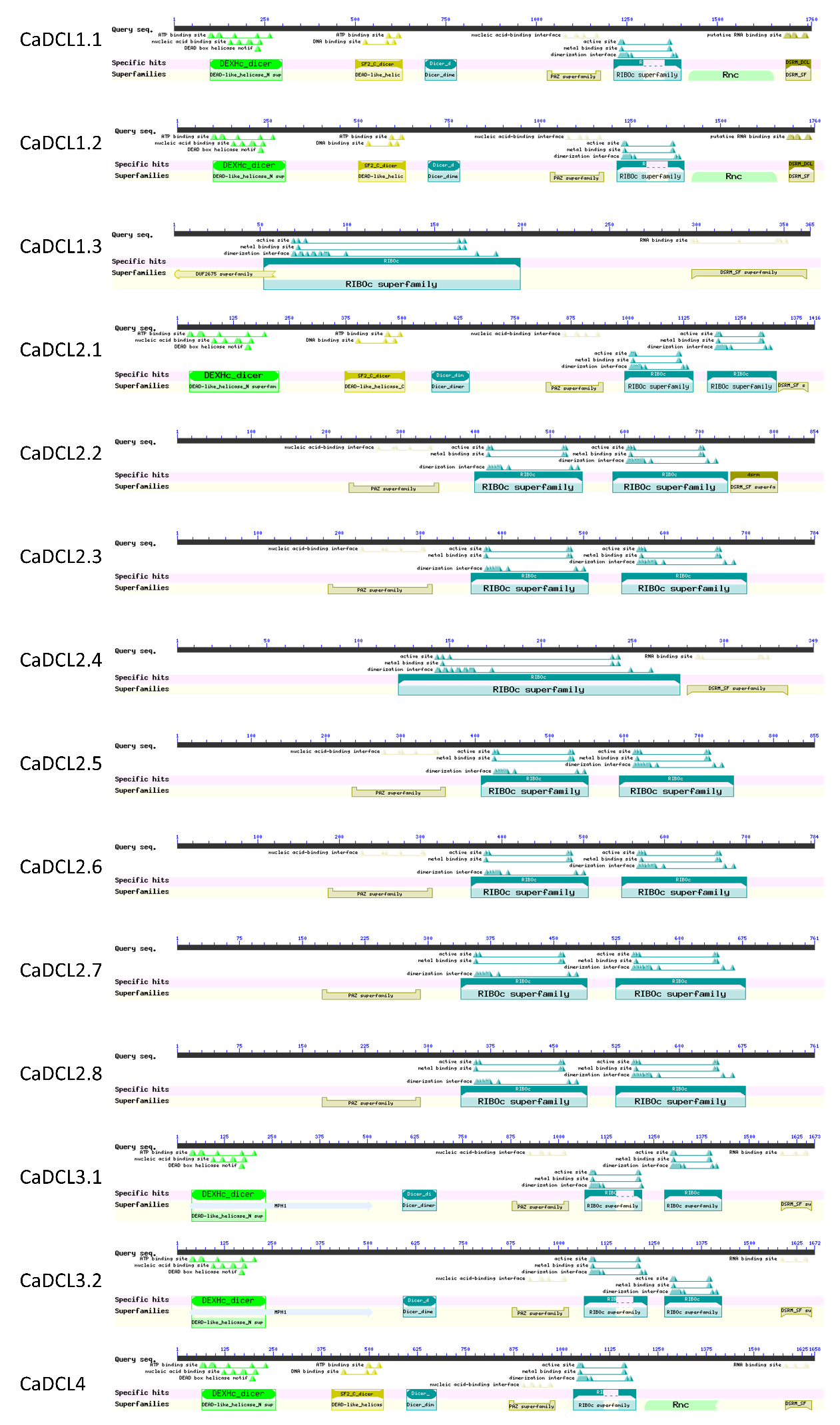

**Fig. S1 - DCL domain architecture from 14 *C. arabica* putative proteins generated by Conserved Domains software** (available at https://www.ncbi.nlm.nih.gov/cdd). Putative proteins predicted presenting Ribonuclease_3, DEAD/RES III, Helicase_C, Dicer_dimer, PAZ, and DSRM domains.

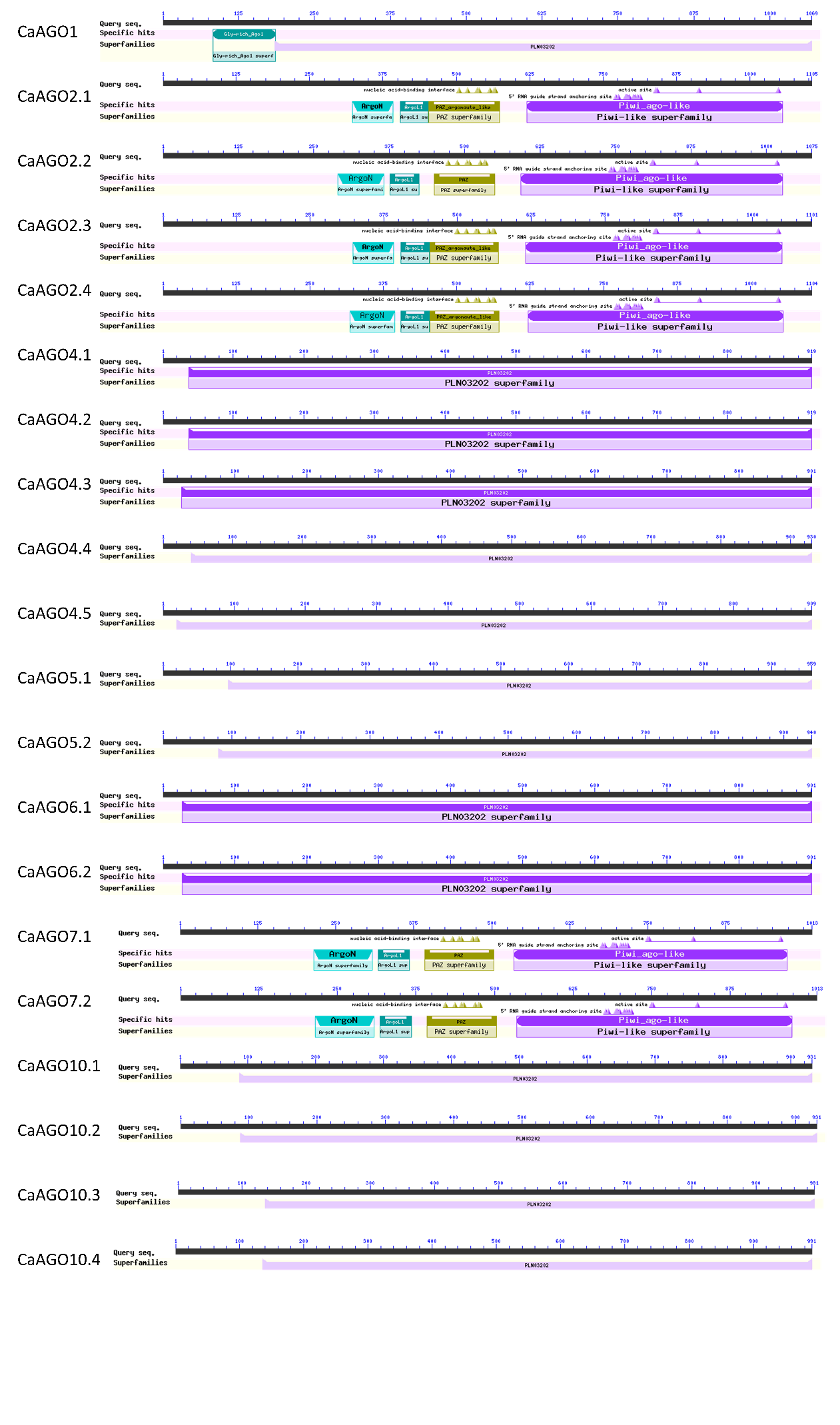

Fig. S2- AGO domain architecture from 20 *C. arabica* putative proteins generated by Conserved Domains software (available at https://www.ncbi.nlm.nih.gov/cdd). Putative proteins predicted presenting ArgoN, PAZ, Piwi, ArgoL1, ArgoL2 and ArgoMID and Gly-rich AGO 1 domains.
